# RBFOX2 is Critical for Maintaining Alternative Polyadenylation Patterns and Mitochondrial Health in Rat Myoblasts

**DOI:** 10.1101/2020.05.13.093013

**Authors:** Jun Cao, Sunil K. Verma, Elizabeth Jaworski, Stephanie Mohan, Chloe K. Nagasawa, Kempaiah Rayavara, Amanda Sooter, Sierra N. Miller, Richard J. Holcomb, Ping Ji, Nathan D. Elrod, Eda Yildirim, Eric J. Wagner, Vsevolod Popov, Nisha J. Garg, Andrew L. Routh, Muge N. Kuyumcu-Martinez

## Abstract

RBFOX2, which has a well-established role in alternative splicing, is linked to heart diseases. However, it is unclear whether RBFOX2 has other roles in RNA processing that can influence gene expression/function in muscle cells, contributing to disease pathology. Here, we employed both 3’-end and nanopore cDNA sequencing to reveal a previously unrecognized role for RBFOX2 in maintaining alternative polyadenylation (APA) signatures in myoblasts. We found that RBFOX2-mediated APA modulates both mRNA levels and isoform expression of a collection of genes including contractile and mitochondrial genes. We identified the key muscle-specific contractile gene, *Tropomyosin 1* and essential mitochondrial gene, *Slc25a4* as APA targets of RBFOX2. Unexpectedly, depletion of RBFOX2 adversely affected mitochondrial health in myoblasts that is in part mediated by disrupted APA of mitochondrial gene *Slc25a4*. Mechanistically, we found that RBFOX2 regulation of *Slc25a4* APA is mediated through consensus RBFOX2 binding motifs near the distal polyadenylation site enforcing the use of the proximal polyadenylation site. In sum, our results unveiled a new role for RBFOX2 in fine tuning expression levels of mitochondrial and contractile genes via APA in myoblasts relevant to heart diseases.

## INTRODUCTION

RBFOX2 belongs to a family of RNA binding proteins including RBFOX1 and RBFOX3 (neuron specific). RBFOX2 has a well-characterized role in alternative splicing (AS) regulation of pre-mRNAs that can affect gene expression and function. RBFOX2 controls AS by binding to a highly conserved motif (U)GCAUG in intronic and/or exonic regions of pre-mRNAs (Huang et al., 2012; Lovci et al., 2013; Sun et al., 2012; Yeo et al., 2009) and regulates AS in a large complex with other splicing regulators (Damianov et al., 2016). RBFOX2 has been linked to congenital heart defects (Homsy et al., 2015; Verma et al., 2016b), heart failure (Wei et al., 2015b), cardiac complications of diabetes (Nutter et al., 2017; Nutter et al., 2016b) and cardiac arrhythmias in Myotonic Dystrophy (Misra et al., 2020). RBFOX2 has been shown to be important for skeletal muscle development and function (Singh et al., 2018; Singh et al., 2014). Due to its important role in AS, the majority of studies have been on identification of RBFOX2-regulated AS events in heart and skeletal muscle (Runfola et al., 2015; Singh et al., 2018; Verma et al., 2016a). However, it remains unclear how loss of RBFOX2 contributes to heart and skeletal muscle defects. Recent studies have suggested that RBFOX2 binds within the 3’UTR of transcripts (Lovci et al., 2013; Wang et al., 2008; Weyn-Vanhentenryck et al., 2014) in proximity with poly(A) sites (PASs) (Wang et al., 2008). The consequences of this binding are unknown and a role for RBFOX2 in cleavage and polyadenylation of pre-mRNAs has not yet been demonstrated.

Mammalian 3’end processing involves the cleavage of the pre-mRNA near the PASs followed by polyadenylation of the cleaved RNA. This process is complex and requires both *cis*-acting elements in the RNA and *trans*-acting proteins that bind to these elements and regulate cleavage and polyadenylation. Greater than 50% of genes in the human genome present more than one PASs and thus can generate multiple mRNA isoforms defined by different 3’ termini (Tian et al., 2005). Alternative polyadenylation (APA) is defined by the differential usage of these PASs in a given mRNA (Shi, 2012). APA can regulate generation of multiple gene isoforms and impact mRNA expression and homoeostasis (Derti et al., 2012). There are two major types of APA: tandem-APA and splicing-APA. Tandem-APA is mediated by the preferential usage of tandem PASs within a 3’UTR. This type of APA can affect 3’UTR length modulating the presence of cis-acting regulatory elements and/or microRNA binding sites that can control mRNA stability, localization or translation (Di Giammartino et al., 2011). Splicing-APA can lead to generation of multiple protein isoforms with distinct functions and properties via AS of the terminal exons that have PASs (Cooke et al., 1999). Tight regulation of APA is essential for normal growth and development and as a result aberrant APA has been linked to many disease states including heart failure, cancer, and muscle disease (Batra et al., 2014; Creemers et al., 2016; Masamha et al., 2014).

In this study, we investigated whether RBFOX2 influences APA decisions in H9c2 myoblasts. We used long-read nanopore cDNA sequencing and Poly(A)-ClickSeq (PAC-seq) (Routh et al., 2017), which allow precise genome wide determination of APA and can quantify gene expression and isoform generation. We identified 233 genes with APA changes upon RBFOX2 depletion in H9c2 myoblasts. 43% of these genes also exhibited mRNA level changes. Further analysis showed that 136 genes that displayed 3’UTR length changes in RBFOX2 depleted myoblasts are known targets of microRNAs, suggesting that modulation of 3’UTR length can impact protein production. We identified RBFOX2 as a regulator of both tandem and splicing-APA, impacting gene expression and isoform generation. Importantly, our results identified mitochondrial gene *Slc25a4* and contractile gene *Tropomyosin 1*, as APA targets of RBFOX2. Investigating how RBFOX2 impacts APA-mediated rat *Slc25a4* expression revealed that RBFOX2 binding motifs located <79nt from the distal polyA site in *Slc25a4* are inhibitory for polyA site usage, negatively modulating *Slc25a4* protein levels. Moreover, the loss of *Slc25a4* APA regulation in RBFOX2-depleted cells correlated with a striking increase in mitochondrial membrane potential and swelling. In summary, we report a new role for RBFOX2 in tuning APA patterns and mitochondrial health.

## RESULTS

### RBFOX2 depletion alters global alternative polyadenylation patterns of pre-mRNAs in H9c2 myoblasts

RBFOX2 binds to the 3’UTR of transcripts (Lovci et al., 2013; Wang et al., 2008; Weyn-Vanhentenryck et al., 2014) near to the PASs (Wang et al., 2008), suggesting a potential role in APA. We have previously shown that H9c2 myoblasts successfully recapitulated RBFOX2-mediated AS regulation observed in mouse hearts (Nutter et al., 2016a), making it a good cell culture model to study RBFOX2 function *in vitro*. Thus, we tested whether RBFOX2 influences APA patterns in rat H9c2 myoblasts. We treated H9c2 cells with scrambled or *Rbfox2*-specific siRNAs and validated KD of RBFOX2 by Western blot (WB) (Fig. 1A). We then isolated RNA from these cells and performed PAC-seq (Routh et al., 2017) and analyzed the data using the differential-poly(A)-clustering (DPAC) pipeline (Routh, 2019a). We detected 101,338 unique PASs with at least 5 unique reads mapping from at least one sample. Multiple PASs occurring within 10nts were clustered together to yield 81,035 unique PASs. We further filtered out all PASs that did not map to 3’UTRs already annotated in the UCSC RefSeq database. After instituting these filters, remaining 22,768 PASs mapped to 10,964 unique mRNA transcripts, of which 5,317 contained multiple PASs and 556 contained multiple terminal exons (TE). DPAC reported 233 APA changes in RBFOX2 depleted myoblasts based on the criteria that differential PAS usage was reported if an individual PAS has a fold change of >1.5, and results with an Independent Hypothesis Weighted (Ignatiadis et al., 2016) multiple testing p-adjusted value of < 0.1 (Fig. 1B, Supp. Data File 1).

**Figure 1.**
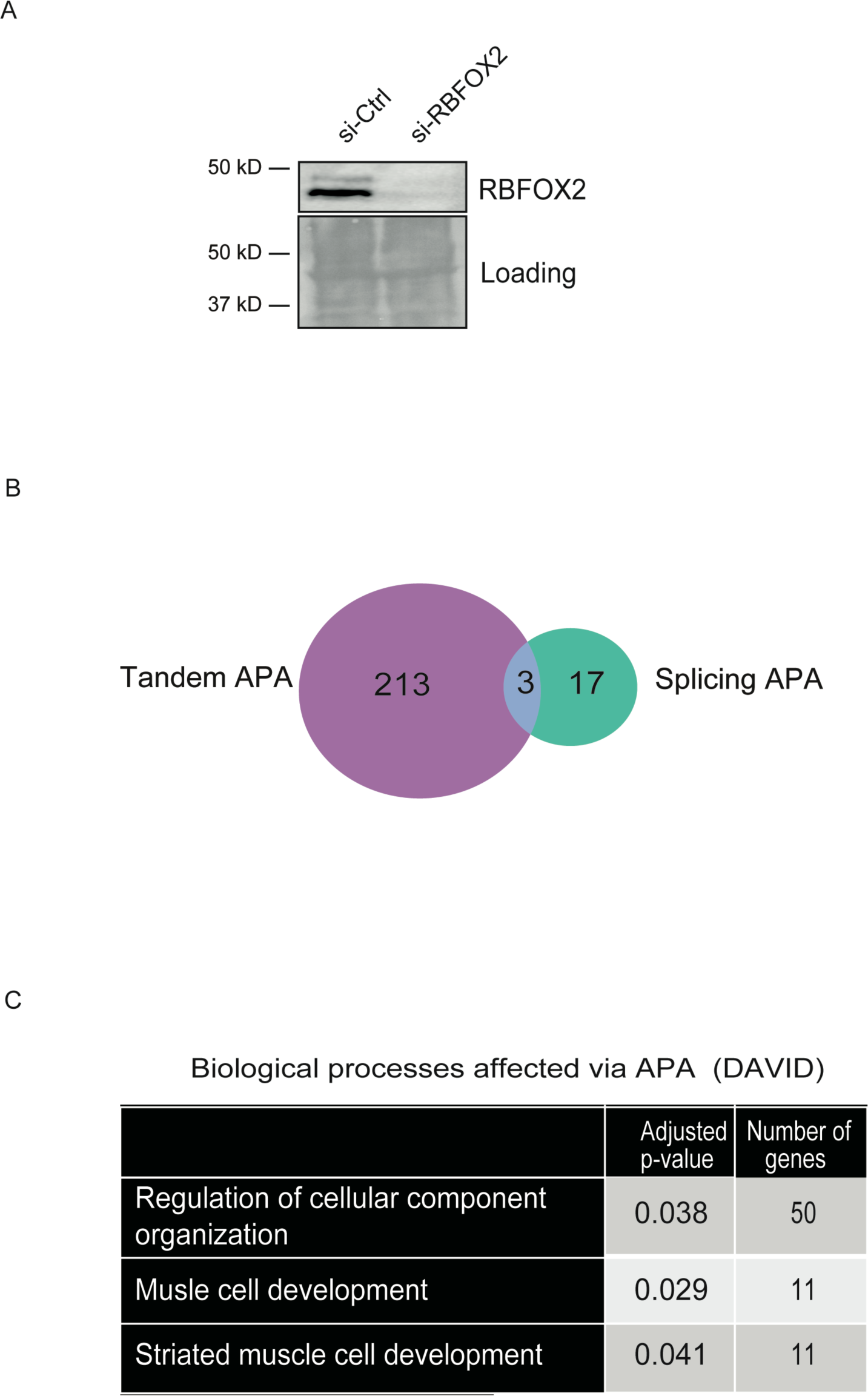
RBFOX2 depletion in H9c2 myoblasts leads to altered alternative polyadenylation patterns determined by Poly(A)-ClickSeq (PAC-seq). **(A)** A representative Western blot showing efficient knock down (KD) of RBFOX2 in H9c2 myoblasts. Ponceau stained membrane was used to monitor protein loading in each lane. **(B)** The number of genes undergo tandem-APA or splicing-APA in RBFOX2 KD H9c2 myoblasts. **(C)** Gene ontology (GO) analysis of genes that undergo APA changes in RBFOX2 depleted myoblasts.

We investigated what types of APA were affected upon RBFOX2 depletion in H9c2 cells. Out of 233 APA events, 213 were tandem-APA, 17 were splicing-APA, and 3 displayed both splicing and tandem-APA (Fig. 1B) (Supp. Data File 2). Using the Gene Ontology (GO) biological function analysis of these APA events, we found an enrichment of genes involved in striated muscle development and regulation of cell organization (Fig. 1C) (Supp. Data File 3). This is consistent with RBFOX2’s involvement in congenital heart defects (Homsy et al., 2015; Verma et al., 2016b) and muscle development (Singh et al., 2018; Singh et al., 2014).

### RBFOX2-mediated splicing-APA is important for expression of rat Tropomyosin 1 muscle-specific isoforms

Based on the GO analysis of RBFOX2 regulated APA events, we first investigated genes that have roles in striated muscle development that are impacted by RBFOX2 mediated splicing-APA (Supp. Data File 2 and Data File 3). Of the 17 splicing-APA changes identified in RBFOX2 depleted myoblasts, *Tropomyosin 1 gene* (*Tpm1*) has an essential role in striated muscle development and function and undergoes a dramatic splicing-APA change in RBFOX2 depleted cells (Supp. Data File 2 and Data File 3). *Tpm1* is an essential gene required for myofibril organization (Thomas et al., 2010), muscle contraction (Wolska and Wieczorek, 2003), and cardiac development (England et al., 2017). *Tpm1* has very complex AS and APA patterns and many isoforms that are tissue specific and developmentally regulated (Gooding and Smith, 2008). Thus, we further studied splicing-APA regulation of rat *Tpm1* by RBFOX2.

PAC-seq identified 2 PASs located in 2 different last exons (exon 9b and exon 9d) that are utilized in H9c2 myoblasts (Fig. 2A). *Tpm1* gene isoforms ending with exon 9b are important for muscle contraction in muscle cells (Lin et al., 2008) (short or muscle-specific transcripts); whereas gene isoforms ending with exon 9d are ubiquitously expressed to support cytoskeleton function (long or non-muscle transcripts). The usage of PAS in muscle specific exon 9b diminished in RBFOX2 knocked down (KD) myoblasts (Fig. 2A).

**Figure 2.**
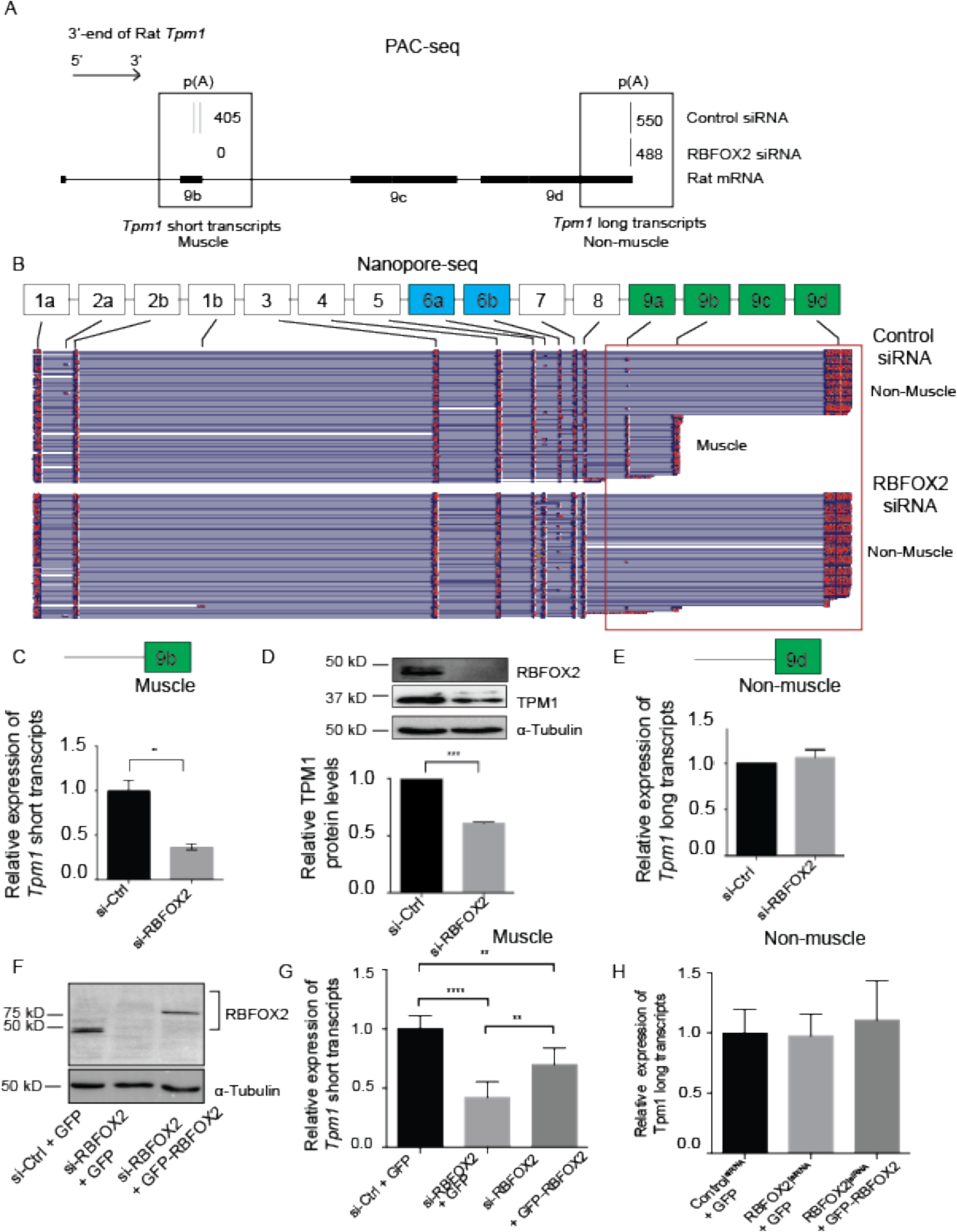
RBFOX2 regulates expression levels of muscle-specific isoforms of rat *Tropomyosin 1* (*Tpm1*) via splicing-APA in H9c2 myoblasts. **(A)** The poly(A) usage of *Tpm1* transcripts in control and RBFOX2 depleted H9c2 cells determined by poly(A)-ClickSeq, (n=3). PolyA sites located in either terminal exon 9b or terminal exon 9d were utilized in H9c2 myoblasts. **(B)** Full-length *Tpm1* short (muscle) or long (non-muscle) isoforms identified by nanopore sequencing in control vs RBFOX2 depleted H9c2 myoblasts (n=3). **(C)** Relative mRNA levels of *Tpm1* muscle-specific (short) transcripts that end with exon 9b in control and RBFOX2 depleted H9c2 cells were determined by RT-qPCR. mRNA levels in control cells were normalized to 1. Data represent means ± SD. Statistical significance was calculated using t-test to compare two different groups in three independent experiments (n=3). **p= 0.0079. **(D)** Relative protein levels of TPM1 in control and RBFOX2 depleted H9c2 cells determined by Western blot. α-Tubulin was used as a loading control. Protein levels in control cells were normalized to 1. Data represent means ± SD. Statistical significance was calculated using t-test to compare two different groups in three independent experiments (n=3) ***p=0.0001. **(E)** Relative mRNA levels of *Tpm1* non-muscle transcripts (long) in control and RBFOX2 depleted H9c2 cells. mRNA levels in control cells were normalized to 1. Data represent means ± SD. No statistical significance was found using t-test in three independent experiments (n=3). **(F)** RBFOX2 protein expression levels in (1) scrambled siRNA treated, (2) RBFOX2 siRNA treated, (3) RBFOX2 siRNA treated H9c2 cells ectopically expressing GFP or GFP-RBFOX2. (1) Ectopic expression of GFP-RBFOX2 partially rescued splicing-APA change in *Tpm1* caused by RBFOX2 depletion. Relative expression levels of muscle-specific *Tpm1* (short) transcripts in scrambled siRNA treated, RBFOX2 siRNA treated, or RBFOX2 siRNA treated H9c2 myoblasts expressing GFP or GFP-RBFOX2. mRNA levels in control cells were normalized to 1. Data represent means ± SD. Statistical significance was calculated using one-way ANOVA to compare three different groups in three independent experiments (n=3). P-value for Control^siRNA^+GFP vs. RBFOX2^siRNA^+GFP is ****p<0.0001; for RBFOX2^siRNA^+ GFP vs. RBFOX2^siRNA^+ GFP-RBFOX2 is **p= 0.0056; for Control^siRNA^+GFP vs. RBFOX2^siRNA^+ GFP-RBFOX2 is **p=0.0027. (2) Relative expression levels of non-muscle *Tpm1* (long) transcripts in control, RBFOX2 depleted, or RBFOX2 depleted GFP or GFP-RBFOX2 expressing cells. mRNA levels in (1) cells were normalized to 1. Data represent means ± SD. Statistical significance was calculated using one-way ANOVA to compare three different groups in three independent experiments (n=3).

To confirm the results obtained by PAC-seq, we used nanopore cDNA sequencing as an orthogonal approach to investigate full length *Tpm1* isoforms and their expression in response to changes in RBFOX2 levels. We performed nanopore cDNA sequencing in control vs RBFOX2 KD cells and obtained high quality and full-length nanopore reads that were mapped to the rn6 genome using Minimap2 (Li, 2018) yielding an average of 550,000 mapped reads spanning 3000-4300 unique mRNAs (Supp. Fig. 1 and Supp. Table 1). Reads were mapped to specific rat *Tpm1* exons (Fig. 2B, black rectangles), identifying 22 different full length *Tpm1* transcripts in control H9c2 myoblasts (Supp. Fig. 2). These *Tpm1* transcripts ended with different terminal exons (exon 9a, exon 9b or exon 9d) in control cells (representative images shown in Fig. 2B). Upon depletion of RBFOX2, we observed a strong reduction in the diversity of *Tpm1* isoforms with only nine different full-length transcripts that ended almost exclusively with exon 9d (Fig. 2B and Supp. Fig. 3), consistent with reduced PAS usage in exon 9b in RBFOX2 KD cells (Fig. 2A). These indicate that RBFOX2 KD reduces expression of muscle-specific isoforms that end with exon 9b.

**Figure 3.**
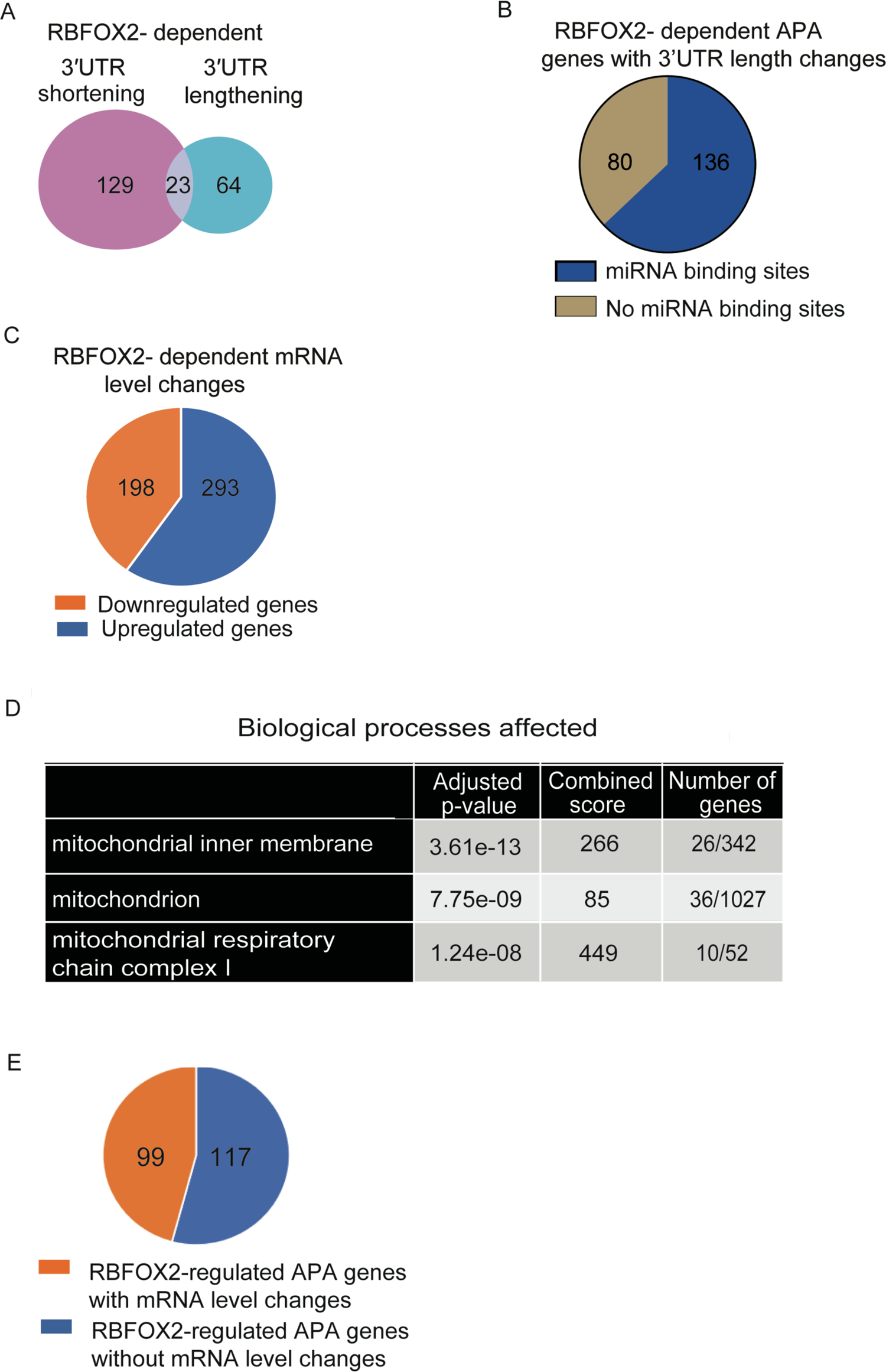
RBFOX2-regulated tandem-APA modulates 3’UTR length and affects mRNA levels. **(A)** The number of genes that undergo tandem-APA changes generating shorter or longer 3’UTRs in RBFOX2 KD H9c2 myoblasts. **(B)** Enrichr TargetScan analysis of genes that display 3’UTR length changes upon RBFOX2 depletion revealed known microRNA binding sites within majority of these genes. **(C)** Gene expression changes in RBFOX2 KD H9c2 myoblasts. **(D)** Gene ontology analysis of genes that are downregulated in RBFOX2 depleted H9c2 myoblasts. **(E)** The number of genes that exhibit both APA and gene expression changes in RBFOX2 KD H9c2 myoblasts.

To further validate APA changes in muscle-specific (short) isoforms of *Tpm1*, we designed primers spanning *Tpm1* exon7 and exon 9b or exon 7 and exon 9d (Supp. Table 2). The levels of *Tpm1* muscle-specific transcripts were dramatically decreased in RBFOX2 depleted cells (Fig. 2C). Consistent with reduced levels of muscle-specific *Tpm1* isoforms, TPM1 protein levels were reduced by 45% in RBFOX2 KD cells (Fig. 2D). Importantly, mRNA levels of *Tpm1* non-muscle (long) transcripts did not change upon RBFOX2 KD (Fig. 2E).

We performed rescue experiments by ectopically expressing GFP-tagged RBFOX2 in RBFOX2 KD H9c2 cells. *Rbfox2* siRNA treatment caused effective KD of RBFOX2 in H9c2 cells, and GFP-RBFOX2 protein was expressed at lower levels than the endogenous RBFOX2 due to targeting by RBFOX2 specific siRNAs (Fig. 2F, lane 3 vs 1). Even low levels of GFP-RBFOX2 partially rescued the expression of muscle-specific *Tpm1* transcripts (Fig. 2G). GFP-RBFOX2 expression did not affect expression of non-muscle *Tpm1* isoforms (Fig. 2H). Importantly, *Tpm1* regulation by RBFOX2 was specific because we did not observe dramatic changes in APA patterns of rat *Tpm2, Tpm3* and *Tpm4* in RBFOX2 KD myoblasts (Supp. Fig. 4).

**Figure 4.**
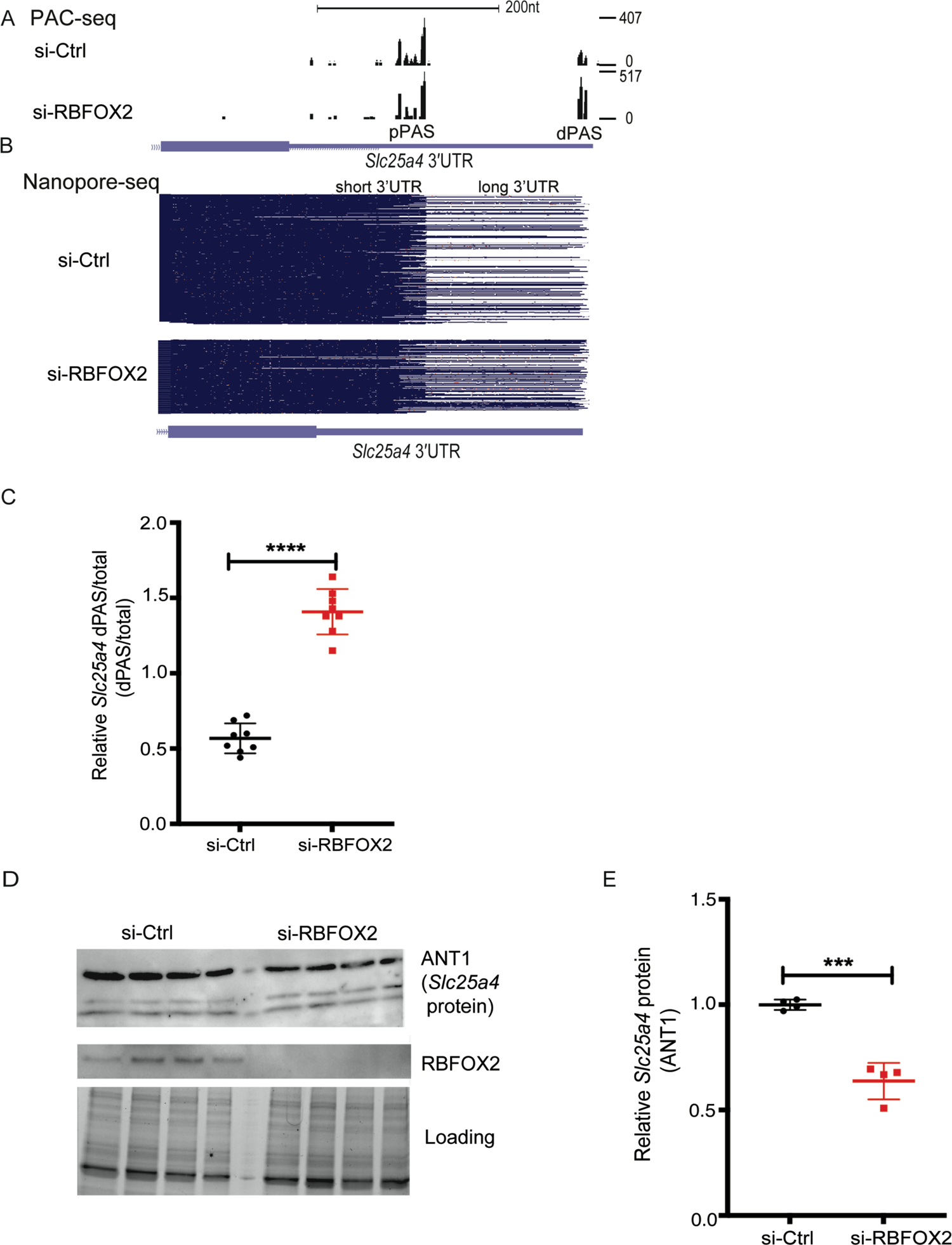
RBFOX2 regulates tandem-APA and expression of essential mitochondrial gene *Slc25a4*. **(A)** *Slc25a4* 3’UTR lengthening mediated via tandem-APA in RBFOX2 depleted H9c2 myoblasts identified by PAC-seq. **(B)** Nanopore sequencing analysis of full length *Slc25a4* transcripts with different 3’UTR lengths in control vs RBFOX2 KD myoblasts. **(C)** RT-PCR analysis of endogenous *Slc25a4* dPAS/total mRNA ratio in RBFOX2 KD myoblasts. n=8, ****p< 0.0001 (three independent experiments). **(D)** Western blot analysis of ANT1 protein (*Slc25a4*) in control vs RBFOX2 KD myoblasts determined by Bio-Rad ChemiDoc Imager. Even protein loading was monitored by imaging stain-free gels. ANT1 protein levels were normalized to loading control and fold change in ANT1 protein levels were quantified using Bio-Rad ChemiDoc software. n=4, ***p= 0.0002.

These data show that RBFOX2 controls expression of muscle-specific isoforms of essential contractile gene *Tpm1* via splicing-APA. Moreover, our results also demonstrate that nanopore sequencing is an effective method in revealing complex and coordinated APA and AS patterns and validating AS and APA variants.

### RBFOX2-mediated tandem-APA alters 3’UTR length and mRNA levels

Having established splicing-APA mediated regulation of *Tpm1* pre-mRNA by RBFOX2, we sought to identify novel tandem-APA events in genes that may be critical for muscle development and function. We found that the majority of the APA changes in RBFOX2 depleted cells were tandem-APA (Fig. 1B), which can modulate the 3’UTR length and, in turn, gene expression levels. Analysis of PAC-seq data revealed that 129 out of 216 (59%) APA changes led to 3’UTR shortening (Supp. Data Files 1 and 4) and 64 out of 216 (29.6%) led to 3’UTR lengthening (Supp. Data Files 1 and 5) in RBFOX2 depleted cells (Fig. 3A). There was a smaller subset that included 23 genes that contained 3 or more PASs giving rise to complex changes resulting in both lengthening and shortening of 3’UTRs in RBFOX2 KD cells due to the modulation of levels of both the most distal and most proximal PASs relative to an intermediate PAS (Fig. 3A) (Supp. Data Files 1 and 6). It has been well-established that 3’UTRs harbor binding sites for microRNAs that can influence mRNA translation and/or stability. We used TargetScan to determine whether genes with 3’UTRs length changes harbor known targets of microRNAs. We found that out of 216 tandem-APA events that display 3’UTR length changes in RBFOX2 KD myoblasts, the majority (136 genes) displayed microRNA binding sites in their 3’UTRs (Fig. 3B) (Supp. Data File 7). These results suggest that changes in 3’UTR length mediated via RBFOX2 depletion could affect microRNA-mediated regulation of these genes.

Given that 3’UTR length can alter mRNA stability, we wondered if genes that exhibit APA changes upon RBFOX2 KD also display expression level changes. There were 491 mRNA level changes (fold Change > 1.5, p-adjust < 0.1) in RBFOX2 KD myoblasts (Fig. 3C) (Supp. Data File 8). GO analysis (Kuleshov et al., 2016) of genes that are downregulated in RBFOX2 KD myoblasts revealed mitochondria, mitochondrial membrane and mitochondrial respiratory chains as affected processes (Fig. 3D) (Supp. Data File 9). There were 26 mitochondrial genes that were downregulated in RBFOX2 depleted myoblasts (Supp. Data File 9), Importantly, our analysis of differential gene expression also revealed that 99 of the 216 (46%) genes with tandem-APA changes in RBFOX2 depleted myoblasts also exhibited changes in mRNA levels (Fig. 3E). Collectively, these data indicate that RBFOX2-mediated regulation of tandem-APA events impact both 3’UTR length and gene expression but also uncovered an enrichment of target genes involved in mitochondrial function.

### RBFOX2-mediated tandem-APA regulates 3’UTR length and protein levels of mitochondrial gene *Slc25a4* that encodes for ATP/ADP translocase 1 (ANT1)

To better understand how RBFOX2 impacts mitochondrial gene expression, we analyzed mitochondrial genes that displayed both tandem-APA and mRNA level changes and are identified in our GO analysis to be important for muscle development (Supp. Data File 3). Our DPAC analysis revealed that total *Slc25a4* mRNA levels are downregulated by 1.63-fold in RBFOX2 depleted cells (Supp. Data File 8). *Slc25a4* gene is the ATP/ADP translocator 1 (ANT1) critical for energy production in the mitochondria. It has been shown that ablation of *Slc25a4* (ANT1) in mice causes cardiomyopathy associated with defects in mitochondrial respiration (Graham et al., 1997). Loss of function mutations in *Slc25a4* are linked to hypertrophic cardiomyopathy and skeletal muscle myopathy in humans (King et al., 2018; Korver-Keularts et al., 2015).

PAC-seq identified a tandem-APA change in *Slc25a4* in RBFOX2 KD cells (Supp. Data File 1). *Slc25a4* has 2 major PASs (distal and proximal) in its 3’UTR (Fig. 5A). While distal PAS (dPAS) was minimally used in control cells, its usage was favored in RBFOX2 KD cells (Fig. 4A). This result from PAC-seq was consistent with the nanopore sequencing data that identified mostly *Slc25a4* transcripts with long 3’UTR generated via dPAS usage in RBFOX2 KD H9c2 cells (Fig. 4B, bottom panel). These results were reproducible in 3 different control and RBFOX2 KD myoblasts (Supp. Fig. 5).

**Figure 5.**
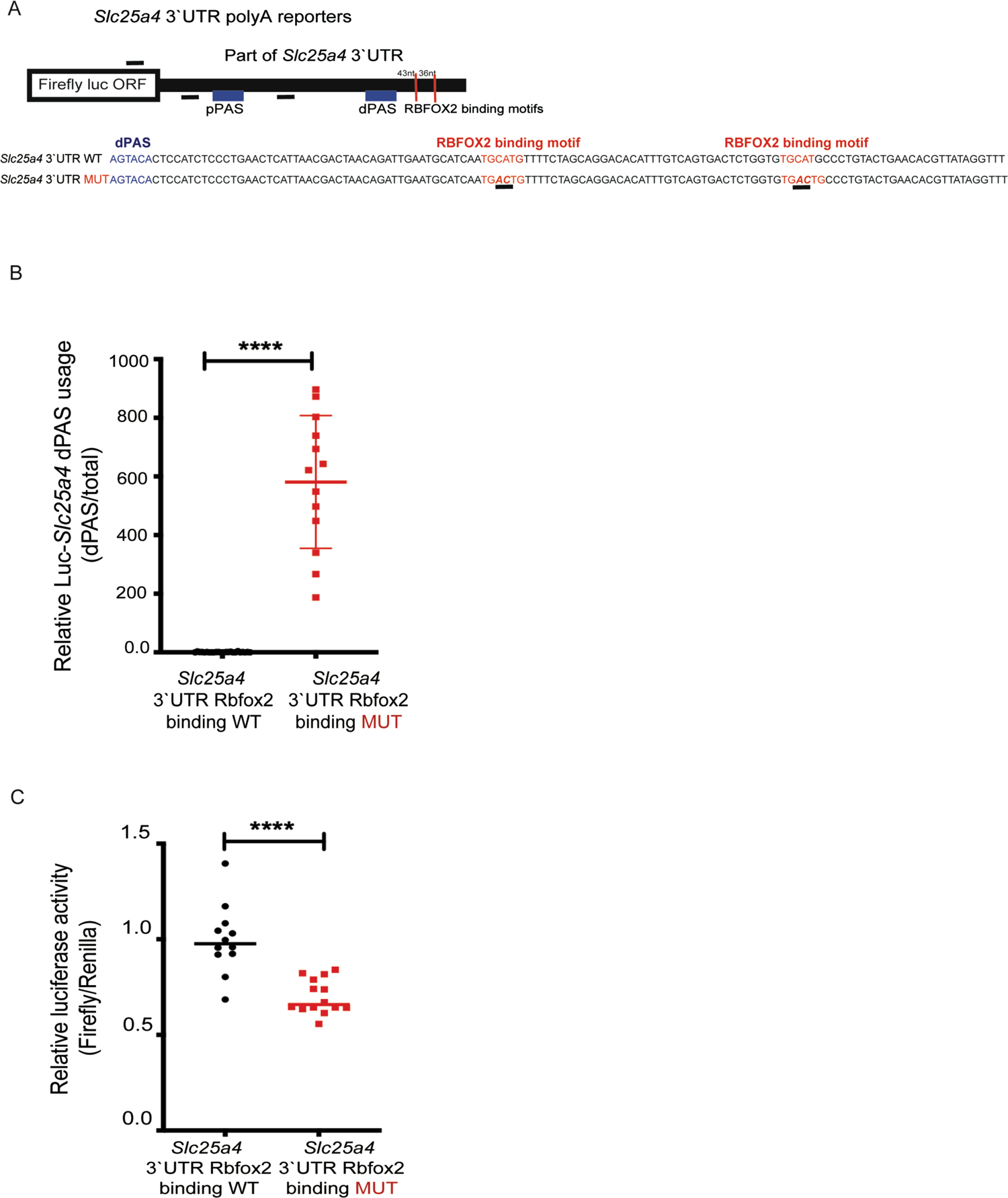
(A) Cartoon representation of Luciferase-*Slc25a4* polyA (pA) reporter constructs that harbor both pPAS and dPAS of *Slc25a4* in the 3’UTR with or without mutated RBFOX2 binding motifs (underlined) located downstream of dPAS. **(B)** RT-qPCR analysis of dPAS usage when normalized to total mRNA levels in HEK293 cells expressing WT or RBFOX2 binding site mutant pA Luciferase-*Slc25a4* 3’UTR constructs, n.6, p< 0.0001 (three independent experiments). **(C)** Relative firefly luciferase levels in HEK293 cells expressing WT or RBFOX2 binding site mutant pA Luciferase-*Slc25a4* 3’UTR constructs, n>5, p< 0.0001 (three independent experiments).

To validate the APA changes in *Slc25a4* transcripts, we designed primers to detect long *Slc25a4* transcripts generated via dPAS usage and total *Slc25a4* transcripts (long and short) (Supp. Table 2). Relative *Slc25a4* dPAS/total mRNA ratio was significantly increased in RBFOX2 KD cells (Fig. 4C), consistent with the PAC-seq and nanopore sequencing data indicating increased dPAS usage (Fig. 4A and 4B). Because increased dPAS usage generates *Slc25a4* transcripts with long 3’UTRs, which can negatively impact mRNA stability and/or translation, we checked *Slc25a4* protein (ANT1) levels. We found that ANT1 protein levels were significantly downregulated in RBFOX2 KD myoblasts (Fig. 4D and 4E), correlating well with the presence of *Slc24a5* transcripts with long 3’UTR (Fig. 4B and 4C). These results indicate that RBFOX2 is critical for *Slc25a4* expression.

### RBFOX2 binding motifs near PAS sites are critical for APA-mediated regulation of *Slc25a4* expression

To better understand how RBFOX2 regulates *Slc25a4* expression via APA, we examined 3’UTR of rat *Slc25a4* and found two RBFOX2 binding consensus binding sites one 43nt and another 79nt away from dPAS of *Slc25a4* (Fig. 5A). To test whether RBFOX2 binding motifs near the dPAS are functionally important for dPAS usage and *Slc25a4* expression, we generated Luciferase-*Slc25a4* 3’UTR polyA heterologous reporters. While generating these constructs we inserted part of the *Slc25a4* 3’UTR that contains pPAS, dPAS and RBFOX2 binding sites, downstream of the firefly luciferase open reading frame. We also removed the SV40 PAS within the plasmid (Fig. 5A). In this way, luciferase mRNA will be cleaved and polyadenylated using the PASs within the *Slc25a4* 3’UTR.

Using this luciferase reporter, we mutated the two RBFOX2 consensus “TGCATG” motifs (Fig. 5A, in red and underlined) near the distal PAS that has been shown to abrogate RBFOX2 binding (Lovci et al., 2013). We tested the relative dPAS usage as well as luciferase activity of the WT construct and compared it to the mutant counterpart that disrupted the RBFOX2 binding site (Fig. 5A, mutant sequence underlined). Mutating RBFOX2 binding motifs in Luciferase-*Scl25a4* 3’UTR polyA reporter resulted in a significant increase in dPAS usage (Fig. 5B, WT vs MUT) and a decrease in firefly luciferase activity (Fig. 5C, WT vs MUT). Firefly luciferase activity was downregulated in RBFOX2 binding site mutants correlating with increased usage of dPAS generating a longer 3’UTR (Fig. 5B), which can reduce mRNA stability and/or translation. These results (Fig. 5B and 5C) are in agreement with the results from PAC-seq, nanopore reads, RT-PCR validations and by Western blot in RBFOX2 depleted myoblasts (Fig. 4). Altogether, these results suggest that RBFOX2 binding sites close to dPAS repress dPAS usage and modulate *Slc25a4* expression levels.

### RBFOX2 depletion adversely impacts mitochondrial health

Since loss of *Slc25a4* (ANT1) is linked to cardiomyopathy and skeletal muscle myopathy and mitochondrial defects and *Slc25a4* (ANT1) protein levels are downregulated in RBFOX2 KD myoblasts, we investigated the status of mitochondria in RBFOX2 depleted cells. Mitochondrial function is essential for heart and muscle function and it declines in human heart failure and diabetic hearts (Rosca and Hoppel, 2013), in which RBFOX2 is implicated. In addition, mitochondrial defects have been identified in hypoplastic left heart syndrome (Karamanlidis et al., 2011; Liu et al., 2017) in which *RBFOX2* mutations have been identified (Homsy et al., 2015; Verma et al., 2016b). The role of RBFOX2 in mitochondrial health is unclear.

Mitochondrial membrane potential is important for respiration and mitochondrion viability (Zorova et al., 2018). This process is affected in RBFOX2 depleted myoblasts (Supp. Data File 9). Therefore, we tested if RBFOX2 KD alters the mitochondrial membrane potential (Ψm), evaluated by the JC-1 assay (Perelman et al., 2012). Mitochondrial membrane potential was increased by 2.5-fold in RBFOX2 KD (vs. control) cells (Fig. 6A). We also used inhibitors of different complexes in the mitochondria to assess mitochondrial membrane changes. In response to rotenone (inhibits complex I), antimycin A (inhibits complex III), FCCP (mitochondrial oxidative phosphorylation uncoupler, depolarizes mitochondrial membrane potential), and H_2_O_2_ treatment, mitochondrial membrane potential was downregulated in control cells as expected. This was also the case in RBFOX2 KD cells; however, RBFOX2 KD cells maintained a higher baseline level of Ψm than was noted in control cells (Fig. 6A). These results may suggest that increased mitochondrial membrane potential upon RBFOX2 deficiency could be to avoid oxidative injury.

**Figure 6.**
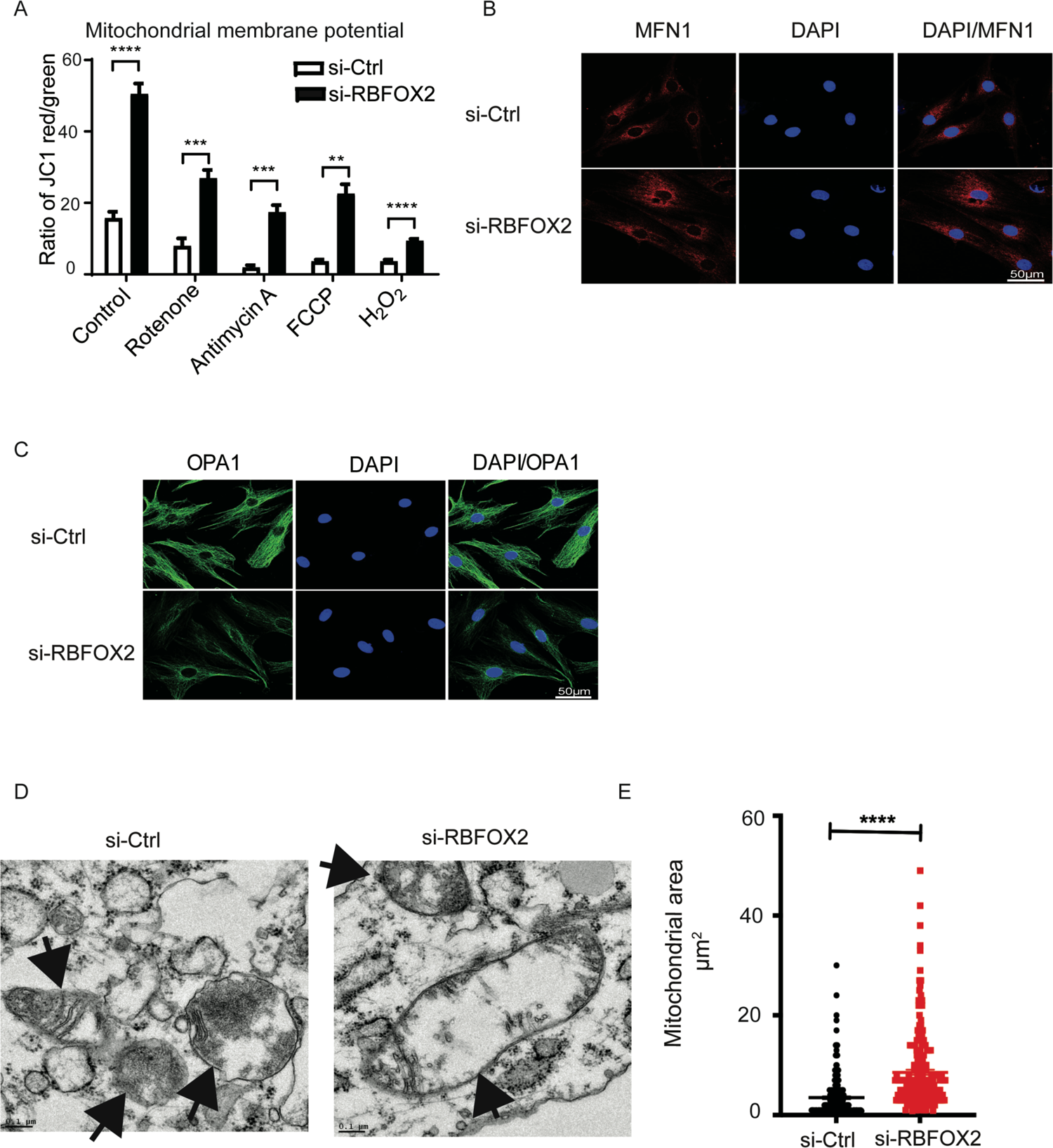
RBFOX2 depletion in H9c2 myoblasts affects mitochondrial health. **(A)** Mitochondrial membrane potential (ψm) was determined by JC-1 staining of the H9c2 cells transfected with scrambled or *Rbfox2* siRNA. Shown are the ratios of fluorescence intensity of J-aggregates (red fluorescence) to J-monomers (green fluorescence). Statistical significance was calculated using unpaired t-test to compare two different groups in five replicates of two independent experiments. Data represent means ± SD. For mitochondrial membrane potential in untreated control vs RBFOX2-KD H9c2 cells, ****p<0.0001; in control vs RBFOX2-KD H9c2 cells treated with Rotenone, ***p= 0.0004; in control vs RBFOX2-KD H9c2 cells treated with Antimycin A, ***p= 0.0009; in control vs RBFOX2-KD H9c2 cells treated with FCCP, **p= 0.0017; in control vs RBFOX2-KD H9c2 cells treated with H_2_O_2_, ****p<0.0001. **(B)** Immunoflourescence of mitochondrial markers OPA1 and **(C)** MFN1 in RBFOX2 KD H9c2 cells imaged by confocal microscopy. **(D)** Representative transmission electron microscopy images of mitochondria in control vs RBFOX2-KD H9c2 cells. Black arrows mark the mitochondria. Scale bar= 0.1 micron **(E)** Surface area of mitochondria in control and RBFOX2 depleted myoblasts was calculated using Image J. (n=234 mitochondria for control, n=216 mitochondria for RBFOX2 KD myoblasts), ****p<0.0001.

Mitochondrial fusion and fission play a key role in maintaining mitochondrial health. Mitofusin 1 (MFN1) is an outer membrane GTPase that mediates mitochondrial clustering and fusion. Dynamin like OPA1 protein regulates mitochondrial fusion and cristae structure in the inner mitochondrial membrane (Song et al., 2009). These two proteins together regulate the Ψm and electron transport chain function and were used as markers to assess electron transport chain function. Our immunofluorescence data showed that RBFOX2 depletion did not overtly impact MFN1 expression but caused a clear decrease in OPA1 levels (Fig. 6B and 6C). Decreased expression of OPA1 protein was consistent with downregulation of its mRNA levels (1.2-fold) in RBFOX2 depleted H9c2 cells (Supp. Data 2). To further determine the effect of RBFOX2 on mitochondrial health, we performed transmission electron microscopy to assess mitochondrial ultrastructure and morphology. We found that in RBFOX2 KD H9c2 cells, mitochondria were swollen as seen in representative micrographs (Fig. 6D, black arrows indicate mitochondria) and also evident from increased mitochondrial area in RBFOX2-depleted cells (Fig. 6E). These type of morphological changes in mitochondria have been linked to the rupture of mitochondria and apoptosis. Our findings indicate that RBFOX2 is important for mitochondrial health, which is relevant to its role in heart diseases.

## DISCUSSION

RBFOX2 is implicated in cardiovascular diseases, congenital heart defects and skeletal muscle development (Gallagher et al., 2011; Homsy et al., 2015; Nutter et al., 2016b; Singh et al., 2018; Singh et al., 2014; Verma et al., 2016b; Wei et al., 2015a). However, it is not clear whether RBFOX2 has additional roles other than AS regulation that influences gene expression and function in muscle cells. In this study, we addressed this fundamentally important question utilizing state of the art techniques and performing functional assays. We uncovered a critical new role for RBFOX2 in maintaining APA patterns in myoblasts and provided insights into the functional consequences of APA changes mediated by RBFOX2 in myoblasts. In addition, our results revealed a novel role for RBFOX2 in regulating mitochondrial health relevant to human heart diseases.

In this study, combined usage of PAC-seq and nanopore sequencing allowed us to precisely determine RBFOX2-regulated APA patterns and evaluate the consequences of RBFOX2-mediated APA changes. PAC-seq allowed us to sensitively detect poly(A)-sites and simultaneously measure transcript abundance, while nanopore sequencing with MinION allowed us to validate PAS usage, simultaneously characterize exon inclusion across the entire mRNA transcript and determine 3’UTR length. By combining PAC-seq and nanopore sequencing, we discovered a new role for RBFOX2 in APA regulation. We identified 233 APA changes in RBFOX2 KD myoblasts and found that 46% of these APA changes impacted mRNA levels. APA regulation is linked to physiological processes including heart development (Nimura et al., 2016), cell proliferation in immune response (Sandberg et al., 2008), and embryonic and postnatal development (Mangone et al., 2010). Aberrant regulation of APA is linked to pathological conditions including heart failure, cancer and muscular dystrophy (Batra et al., 2014; Creemers et al., 2016; Masamha et al., 2014). Here, we showed that RBFOX2-dependent APA events impacted essential contractile gene *Tropomyosin 1*. By identifying developmentally regulated APA networks that impact muscle-specific gene expression and gene isoforms, we provide insights into the post-transcriptional regulation of these genes in the developing muscle.

AS can directly affect PAS usage and APA by regulating the splicing of exons that harbor PASs. RBFOX2 is a well-known regulator of AS. Despite its prominent role in AS, the majority of RBFOX2 induced APA changes were not mediated via splicing-APA. Instead, it was mediated via tandem-APA, which affected the length of 3’UTRs. We validated RBFOX2-mediated splicing-APA and tandem-APA using several different methods including PAC-seq, nanopore sequencing and RT-PCR. We identified rat *Slc25a4* gene as a target of RBFOX2 and found that RBFOX2 binding sites near PASs are important determinants of Slc25a4 dPAS usage and its expression levels. Mutating RBFOX2 binding motifs close to the dPAS increased dPAS usage, suggesting that RBFOX2 may act as a repressor of PAS usage when bound nearby.

In this study, we also focused on the functional consequences of RBFOX2-mediated APA and gene expression changes. We found that RBFOX2 is critical in maintaining mitochondrial gene expression (i.e OPA1 and *Slc25a4*) and mitochondrial health. Importantly, we identified *Slc25a4* gene which encodes for ANT1 protein as a new APA target of RBFOX2. ANT1 is necessary for ATP/ADP translocation across the mitochondrial membrane during oxidative phosphorylation (King et al., 2018; Korver-Keularts et al., 2015). Mutations in this gene is linked to cardiomyopathy in human patients (King et al., 2018; Korver-Keularts et al., 2015). We found that RBFOX2 depletion affected *Slc25a*4 APA, and its expression levels. Consistently, both *Rbfox2* deletion in cardiomyocytes and loss of *Slc25a*4 function resulted in dilated cardiomyopathy (Wei et al., 2015a) (King et al., 2018; Korver-Keularts et al., 2015). Mitochondrial membrane potential is important for ATP production, which is required for heart and muscle function and reduced mitochondrial function is a hallmark of failing hearts (Rosca and Hoppel, 2013). Importantly, we found profound changes in mitochondrial health upon RBFOX2 depletion. Our findings demonstrate that RBFOX2 controls mitochondrial gene expression via APA, providing novel insights into how RBFOX2 may alter mitochondrial health in diseased hearts.

## ACCESSION NUMBERS

All raw sequencing data generated by Nanopore sequencing and PAC-Seq in this manuscript are deposited into the NCBI SRA database with project number PRJNA517125.

## ACKNOWLEDGEMENTS

The authors acknowledge the University of Texas Medical Branch Next Generation Sequencing Core Facility for providing RNA sequencing services. We thank Dr. Pei-Yong Shi and Dr.Ricardo Rajsbaum for allowing us to use their plate readers.

## AUTHOR CONTRIBUTIONS

M.K.M conceived the idea, conceptualized the project, interpreted the data, provided financial support and wrote the manuscript. A.R designed the DPAC pipeline, processed and analyzed the PAC-seq and nanopore data, provided financial support and critically edited the manuscript drafts. J.C conceptualized the project, designed and performed most of the experiments, analyzed data, wrote the manuscript and confirmed the accuracy of data presented in the manuscript. S.K.V. performed experiments related to *Slc25a4* APA, graphed the plots, performed statistics, analyzed the data and provided feedback. S.M. (S. Mohan) performed the analysis for endogenous *Slc25a4* mRNA and protein levels. S.N.M and A.S helped to clone polyA reporters. C.N performed some of the *Slc25a4* RT-PCRs. R.J.H. designed and optimized primers and performed some of the cDNA synthesis. E.J. performed the nanopore sequencing. P.L generated some of the PAC-seq libraries. N.E helped with the preliminary analysis of the PAC-seq data. K.R designed and performed all experiments and analyzed data related to mitochondrial studies. V.P prepared the samples and processed them for TEM and helped analyze the data. E.J.W critically analyzed/interpreted the data and provided feedback for the design of experiments and edited the manuscript drafts. E.Y. provided feedback for the organization of the manuscript and design of the experiments. N.J.G provided financial support for carrying out the mitochondrial function related experiments and helped write and edit the manuscript drafts. All authors have read and approved the manuscript.

## Non-standard Abbreviations and Acronyms

AS: alternative splicing

PAC-seq: poly(A)-ClickSeq

APA: alternative polyadenylation

KD: knock down

3’UTR: 3’untranslated region

PAC: poly(A)-cluster

DPAC: differential-poly(A)-clustering

PAS: Poly(A) site

dPAS: distal poly(A) site

pPAS: proximal poly(A) site

ANT1: (*Slc25a4*) ATP/ADP translocator (mitochondrial) protein

MFN1: Mitofusin

## SUPPLEMENTAL INFORMATION

**Supplemental Figure 1.**
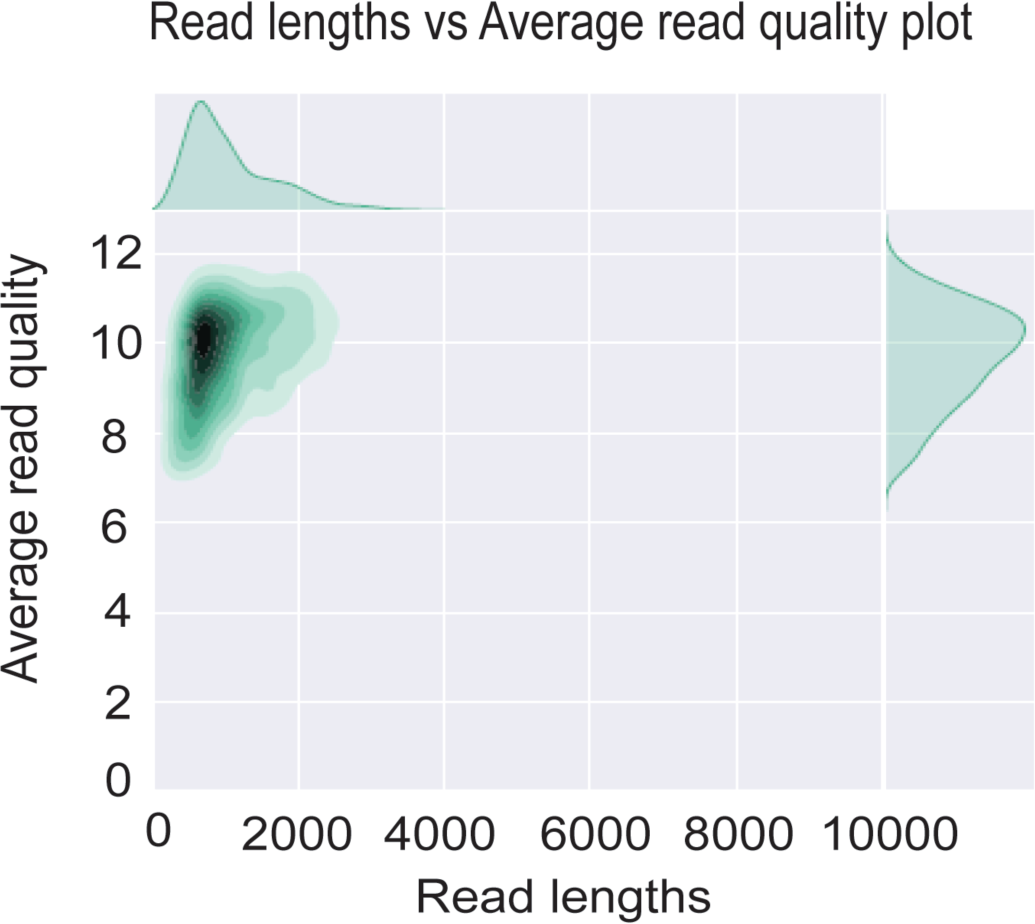
Average read lengths vs read quality of nanopore cDNA sequencing in H9c2 myoblasts.

**Supplemental Figure 2.**
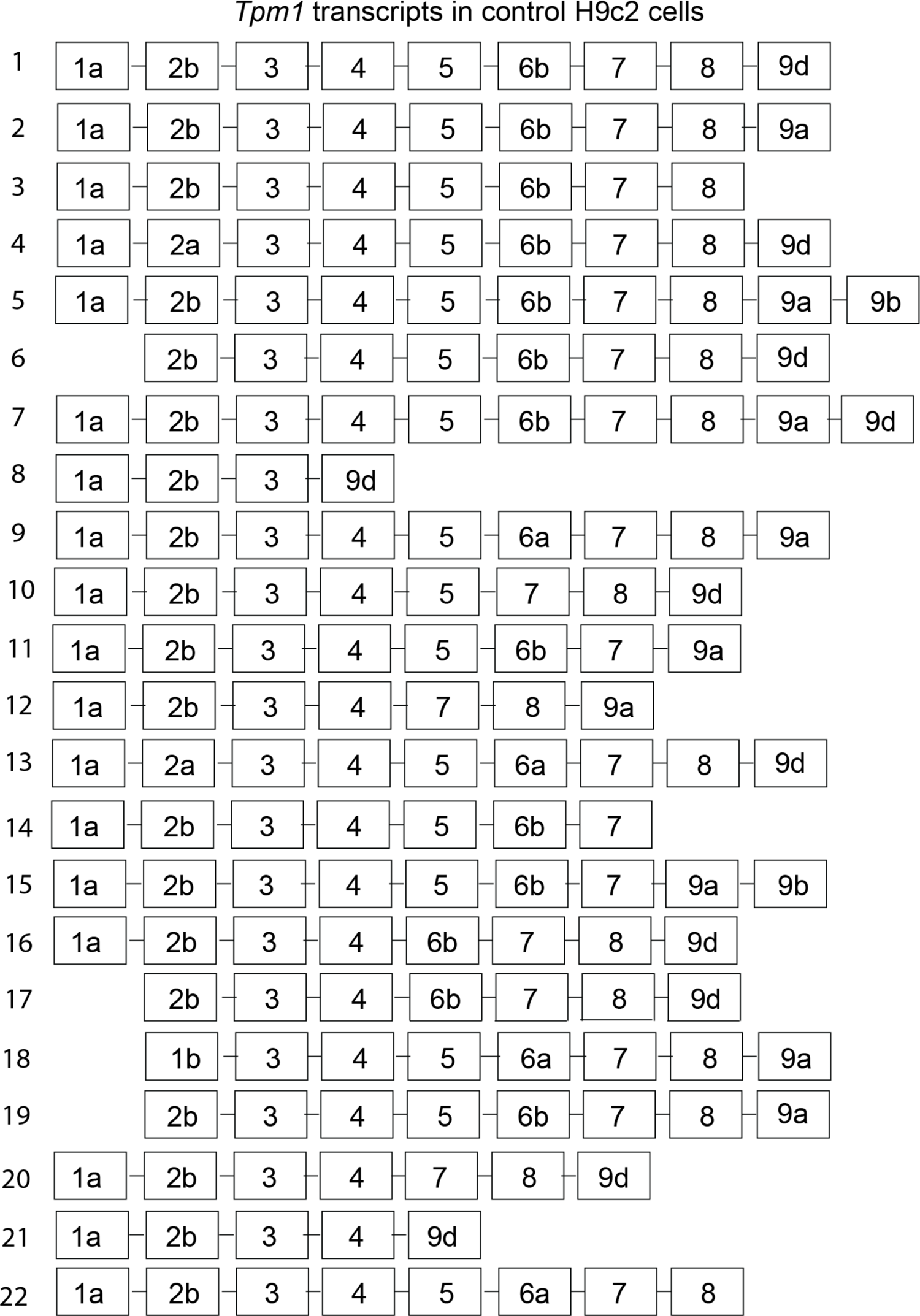
Full-length *Tpm1* transcripts identified by nanopore cDNA sequencing in H9c2 cells.

**Supplemental Figure 3.**
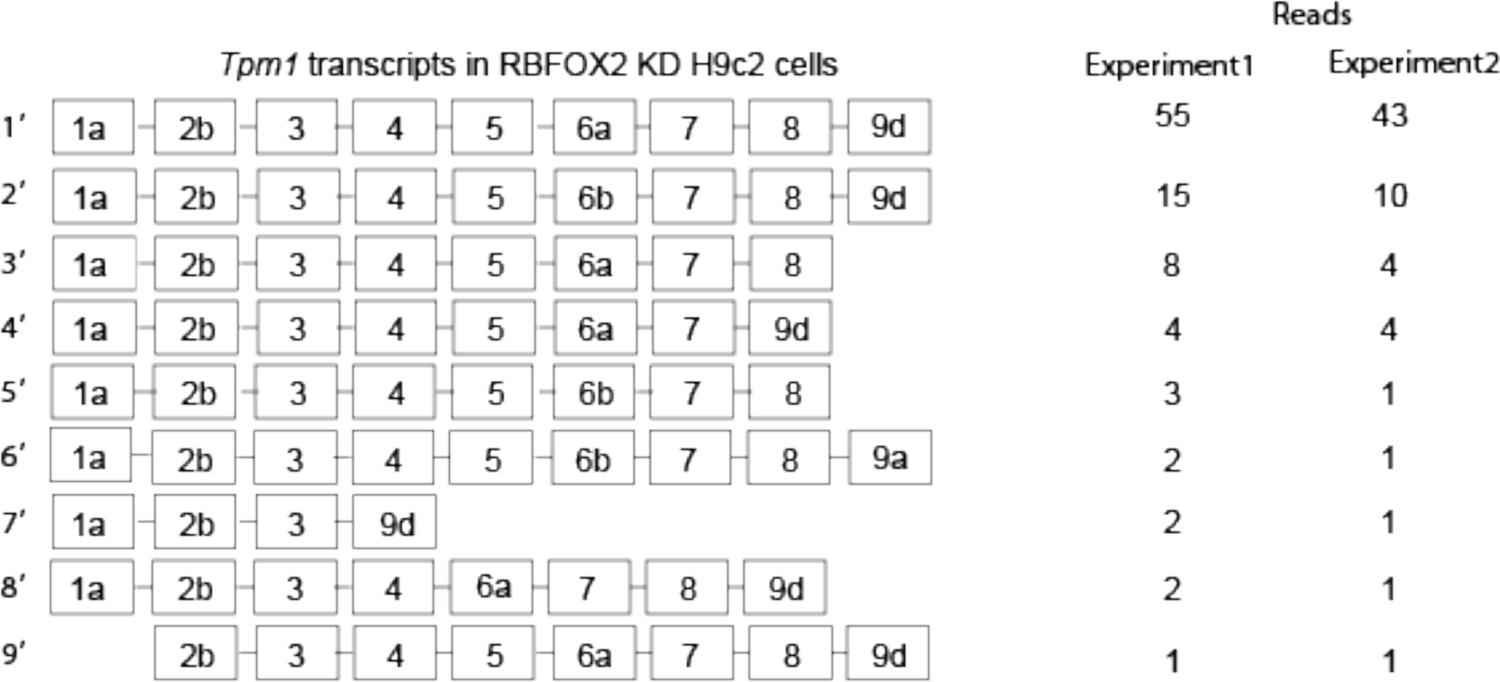
Full-length *Tpm1* transcripts identified by nanopore cDNA sequencing in RBFOX2 knocked down H9c2 cells.

**Supplemental Figure 4.**
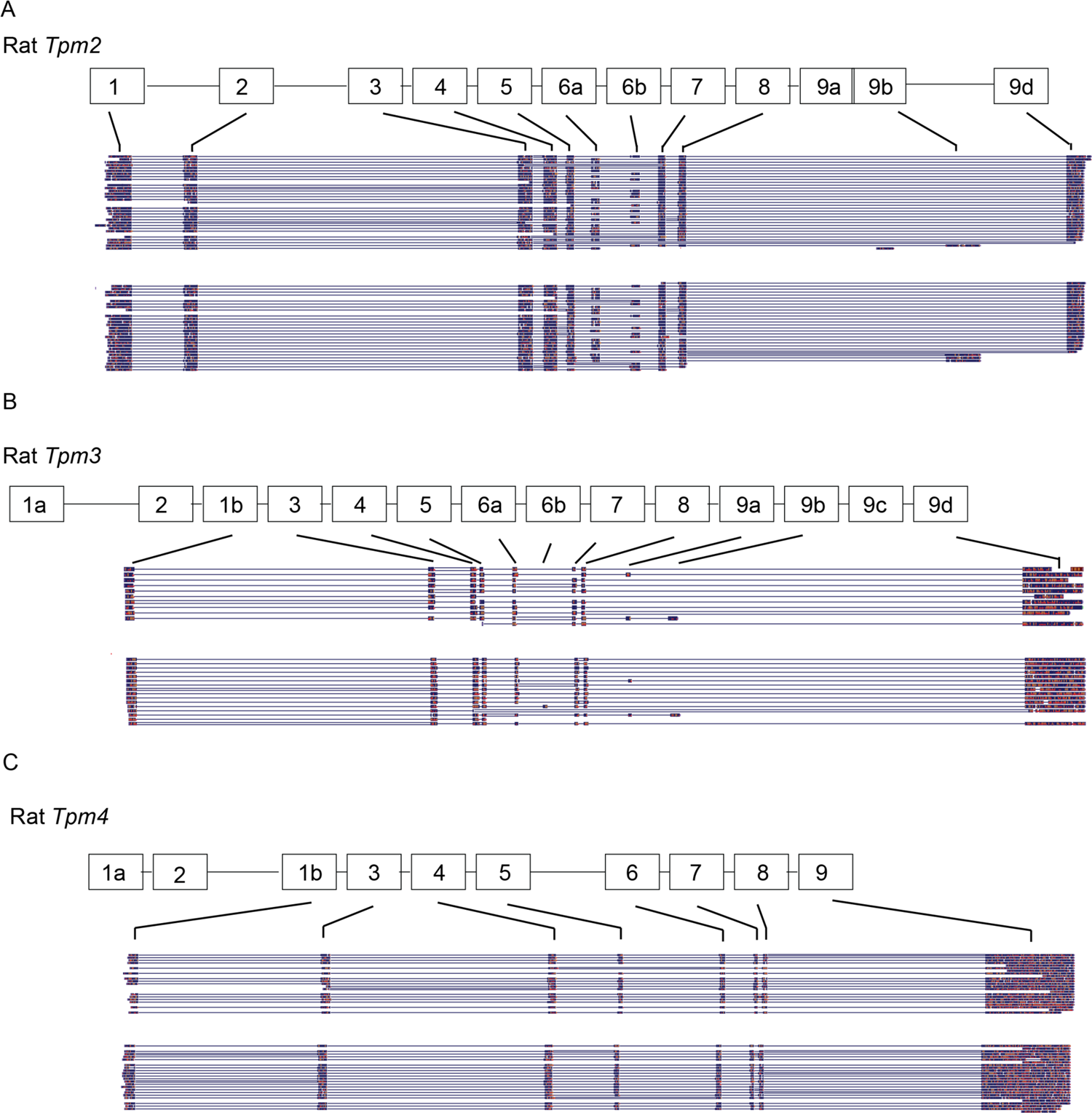
Full length *Tpm2*, *Tpm3*, and *Tpm4* transcripts do not display APA changes in RBFOX2 depleted H9c2 cells determined by nanopore sequencing. **(A)** *Tpm2* transcript sequences mapped to the rat genome after nanopore sequencing in control and RBFOX2 depleted cells. **(B)** *Tpm3* transcript sequences mapped to the rat genome after nanopore sequencing in control and RBFOX2 depleted cells. **(C)** *Tpm4* transcript sequences mapped to the rat genome after nanopore sequencing in control and RBFOX2 KD cells.

**Supplemental Figure 5.**
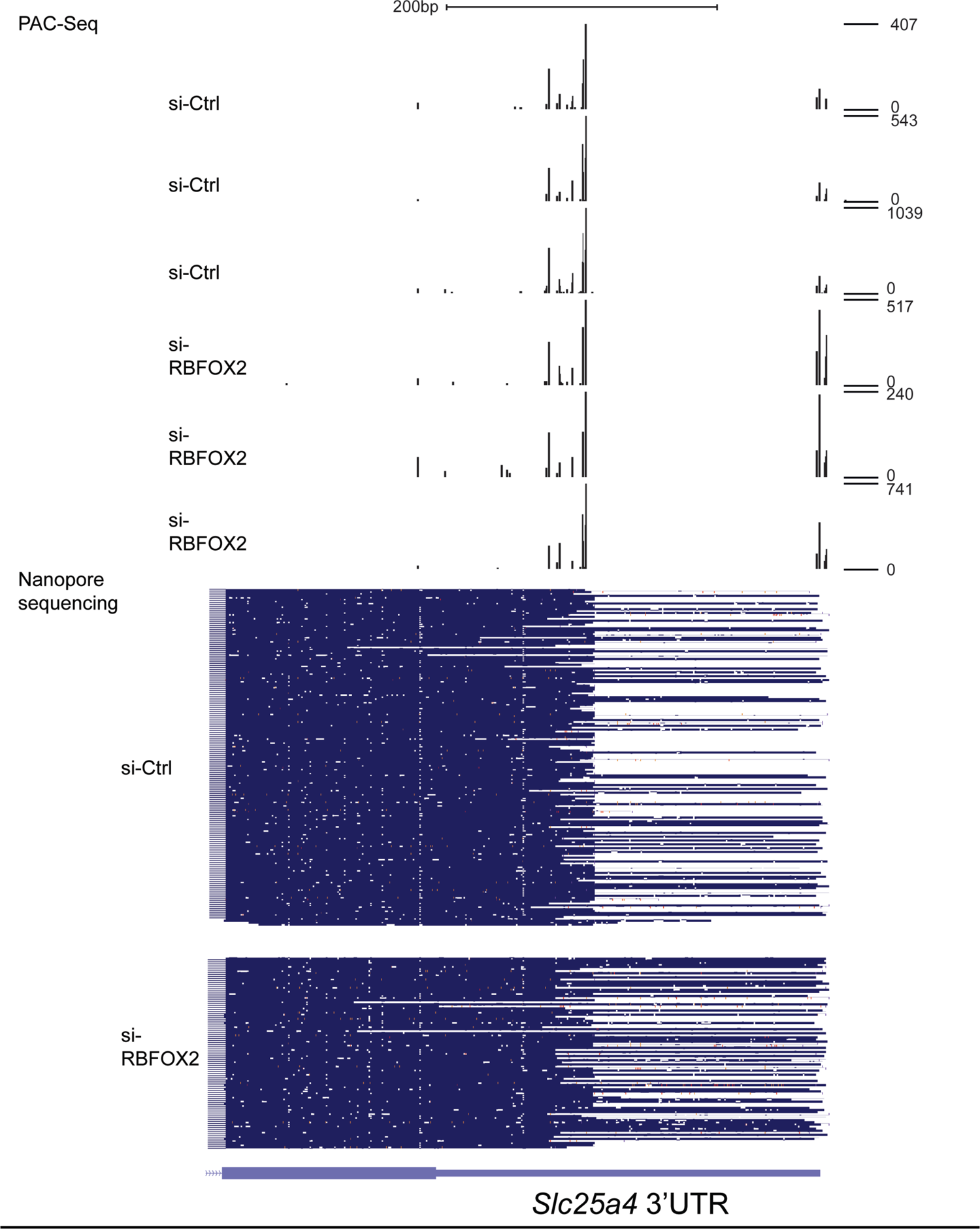
*Slc25a4* PAC-seq and nanopore sequencing data obtained from 3 different control and 3 different RBFOX2 KD H9c2 myoblasts.

## Supplemental Tables

**Supplemental Table 1:**
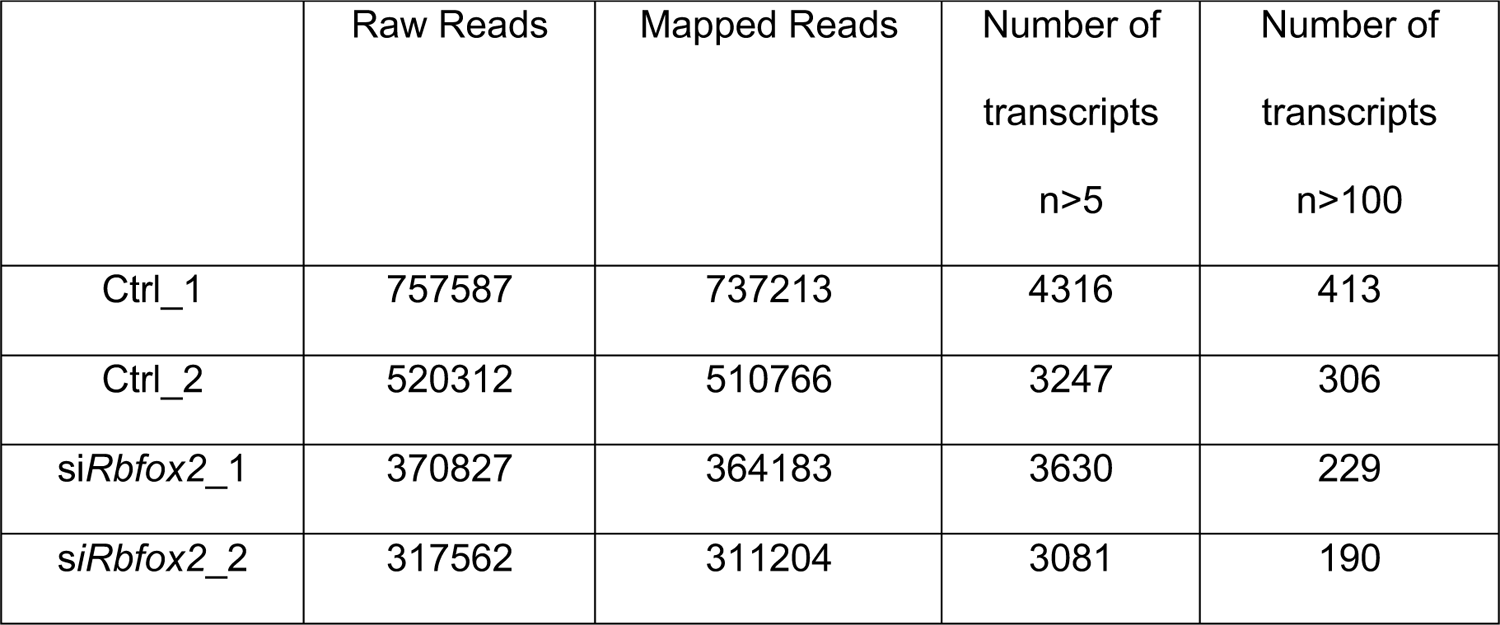
Mapped reads and number of transcripts determined by nanopore sequencing after mapping to rn6 genome using Minimap2.

|  | Raw Reads | Mapped Reads | Number of transcripts<br>n>5 | Number of transcripts<br>n>100 |
| --- | --- | --- | --- | --- |
| Ctrl_1 | 757587 | 737213 | 4316 | 413 |
| Ctrl_2 | 520312 | 510766 | 3247 | 306 |
| siRbfox2_1 | 370827 | 364183 | 3630 | 229 |
| siRbfox2_2 | 317562 | 311204 | 3081 | 190 |

**Supplemental Table 2:**
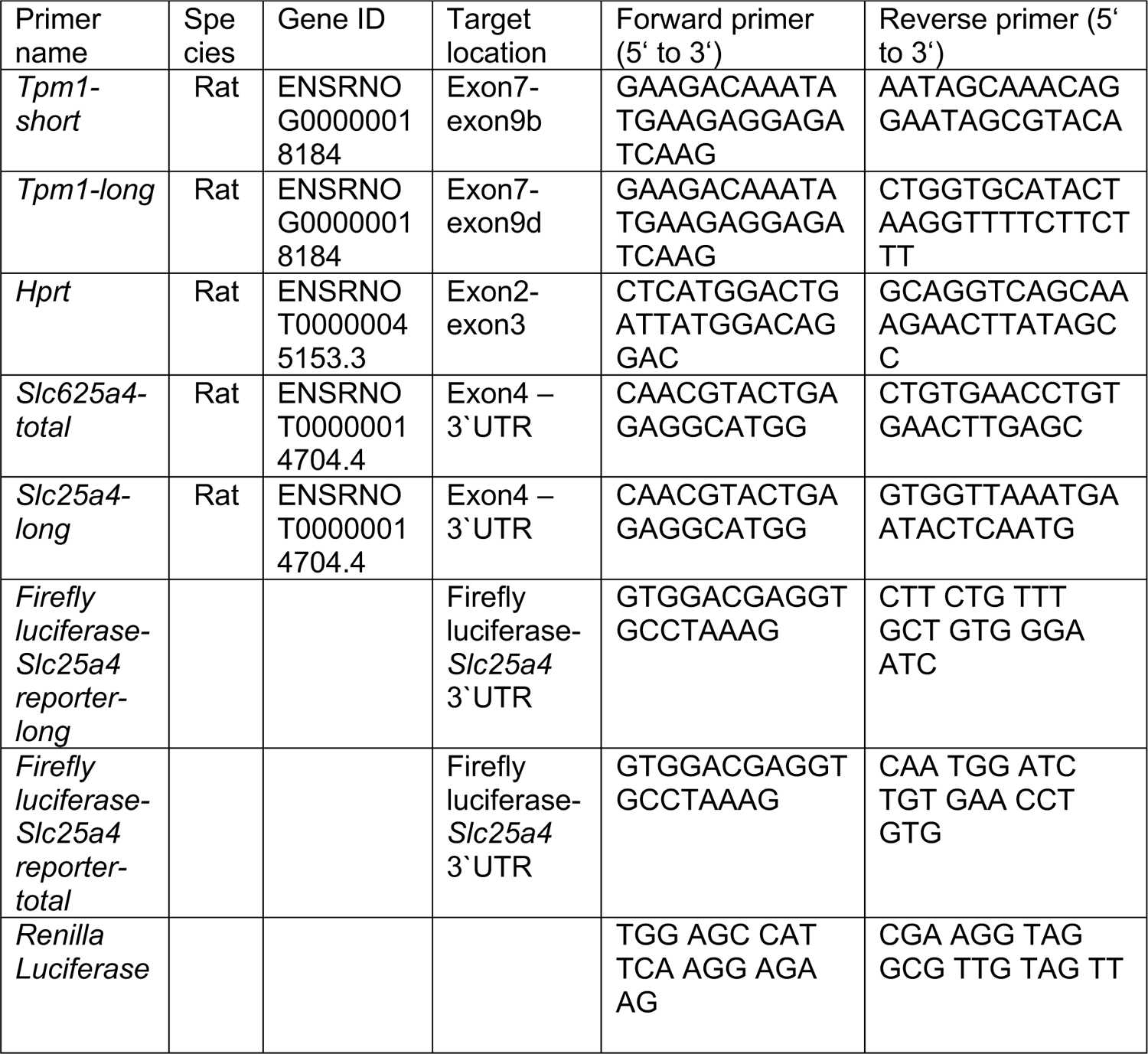
Primer information

## STAR METHODS

### Cell culture

H9c2 myoblast cells (ATCC CRL-1446) were cultured and maintained in Dulbecco’s modified Eagle’s medium (DMEM) (ATCC 30-2002), supplemented with 10% fetal bovine serum (FBS, ATCC 30-2020) and 100 units/ml penicillin and streptomycin (Thermofisher Scientific 15140122). HEK293 cells were maintained in DMEM (Corning 10-013) supplemented with 10% FBS (Corning 35-075), and 100 units/ml penicillin and streptomycin (Thermofisher Scientific 15140122). For siRNA-KD experiments, H9c2 cells were seeded at 10^6^ cells per 100mm dish and transfected with scrambled siRNA (Thermofisher Scientific AM4611) or *Rbfox2* siRNA (Thermofisher Scientific siRNA ID# s96620) at 20nM using Lipofectamine RNAiMAX (Thermofisher Scientific 13778150). Cells were harvested 72 hours post-transfection for RNA or protein extraction. For rescue experiments, 3X10^6^ H9c2 cells were transfected with eGFP (Sigma-Aldrich), human GFP-RBFOX2 (transcript variant 3) (Addgene, plasmid #63086) or empty vector (pcDNA 5) together with scrambled or *Rbfox2* specific siRNAs by Neon Nucleofection System (Thermofisher Scientific) and harvested 48 hours post-transfection as we have described (Verma et al., 2013).

### RNA

RNA was extracted from cells using TRIzol (Thermofisher Scientific 15596-018) by following the manufacturer’s protocol with the exception that RNA was precipitated overnight at −70°C. For PAC-seq, the amount and quality of RNA were analyzed by using an Agilent bioanalyzer at the University of Texas Medical Branch Next-Generation Sequencing Core Facility. For nanopore sequencing, poly(A)+ RNA was enriched by using magnetic mRNA isolation kit (New England biolabs S1550S).

### Poly(A)-ClickSeq (PAC-seq)

The PAC-seq protocol is previously described by Routh et al (Routh et al., 2017). Three samples per treatment/biological condition were used for the PAC-seq analysis. Briefly, 2 µg of total RNA was used to synthesize cDNA through standard reverse transcription initiated by Illumina 4N_21T primer p7 and terminated by incorporation of azido-nucleotides (AzVTPs). Subsequently, the cDNA was cleaned by addition of RNase H (NEB) and purified through DNA Clean and Concentrator Kit (Zymo) before click-reaction, in which the 5’ hexynyl-“click-adapter” p5 was click-ligated with azido-terminated cDNA in the presence of copper-TBTA (Lumiprobe) and Vitamin C. After DNA purification, the clicked cDNA was mixed with indexing primer, universal primer complementary to p7 and One Taq Standard Buffer Master Mix for PCR amplification. PAC-seq libraries were submitted for 1×150 single end sequencing yielding between 14 and 25 million reads per sample. The output data was processed and mapped to the *rattus norvegicus* genome (rn6) and poly(A)-clusters (PACs) were generated using the DPAC pipeline (Routh, 2019b), designed for automated analysis and annotation of poly(A)-targeted RNAseq libraries such as PAC-seq.

*DPAC* (Routh, 2019b) uses DESeq2 (Love et al., 2014) to measure changes in PAC usage between control and RBFOX2 KD cells. Data processing, read mapping and poly(A)-site detection were performed using the default parameters of the *Differential-Poly(A)-Clustering* (*DPAC)* pipeline (Routh, 2019a). PASs are defined as the exact nucleotide of the 3’UTR to poly(A) tract junction found within individual sequencing reads, while poly(A)-clusters (PACs) are defined as clusters of PASs when they are found within <10nts of one another (default parameter in *DPAC* pipeline). The differential PAC usage was reported if an individual PAC comprises at least 5% of a gene’s PACs, has a fold change of >1.5, and results with an Independent Hypothesis Weighted (Ignatiadis et al., 2016) multiple testing p-adjusted value of < 0.1.

### Nanopore sequencing with MinION

For nanopore sequencing, two sets of control and RBFOX2 KD H9c2 cells were used. Total cellular RNA was first poly(A) enriched and then amplified using oligo-dT primers and template switching oligos using Oxford Nanopore Technologies (ONT) cDNA-PCR sequencing kit (PCS108) as prescribed by the manufacturer. Samples were multiplexed using ONT barcodes. Pooled samples were sequenced on R9.4 flow cells. Reads were demultiplexed and base-called using Albacore followed by mapping to the rat genome (rn6) using the splice function of minimap2 (Li, 2016). Transcript isoforms were identified using FLAIR (Tang et al., 2018).

### JC-1 Mitochondrial membrane potential (**Ψm**) assay

H9c2 cells were seeded on to 96-well black flat bottom plates (Costar). After 24 hours, cells were incubated in DMEM media without FBS and phenol red at 37 °C / 5% CO_2_ with 5 µM rotenone (inhibits complex I) or 30 µM antimycin A (inhibits complex III) for 24 hours. In some experiments, cells were also incubated with 250 µM trifluoromethoxy carbonylcyanide phenylhydrazone (FCCP, uncoupler of membrane permeability and oxidative phosphorylation) or 400 µM H_2_O_2_ that was added during last 4 hours of incubation. Plates were washed with PBS and then 100 µl of 10 µg/ml JC-1 (5,5′,6,6′-tetrachloro-1,1′,3,3′-tetraethylbenzimidazolocarbocyanine iodide, Invitrogen, T3168) added. Cells were incubated at 37 °C for 15 minutes in dark, washed twice with PBS, and 100 µl PBS was added to each well. Plates were read in SpectraMax M2 (Molecular Devices) to measure red J-aggregates fluorescence (a sensitive marker of Ψm) at an excitation of 535 nm and emission of 595 nm and green J-monomers fluorescence (indicator of disruption of Ψm) at an excitation of 485 nm and emission of 535 nm. Ratio of red/green fluorescence was calculated and plotted on the graph.

### RT-qPCR

Briefly, a master mix was set up by mixing 5 μl of cDNA, 3 μl of H_2_O, 2 μl of PCR gene specific primer (10X conc) and 10 μl of master mix (Roche 04707516001) in 20 μl reaction. The qRT-PCR was conducted using LightCycler 480 Instrument (Roche) using the following conditions: 95 °C 10 s; 62 °C 15 s; 72 °C 10s for 40 cycles. Melting curve was obtained to ensure single product. ΔCt method was adopted for quantification. qPCR quantifications were as described previously (Belanger et al., 2018; Nutter et al., 2016b; Verma et al., 2016b). For luciferase-*Slc25a4* reporter qPCRs, forward primer was designed on the firefly luciferase ORF, and reverse primers were designed to bind *Slc25a4* 3’UTR in different locations (Supp. Table 2). Firefly-*Slc25a4* heterologous mRNA generated via dPAS usage in comparison to total Firefly-*Slc25a4* heterologous mRNA was normalized to renilla luciferase mRNA levels. Semi-quantitative RT-PCR was used for determining *Tpm1* (short and long) transcripts due to the generation of 2 distinct DNA bands representing short and long isoforms from the same PCR reaction. 2μg of total RNA was used for cDNA synthesis using AMV reverse transcriptase (15 units/μg, Life Biosciences). PCR was performed using 5μl of cDNA, 25μM dNTPs, 100ng of each gene specific forward and reverse primer and 0.2μl of Biolase Taq polymerase (Bioline) in a 20μl reaction. All primer sequences are provided in Supp. Table 2.

### Western blot

Cells were lysed in lysis buffer (10mM HEPES-KOH, pH7.5, 0.32 M Sucrose, 5 µM MG132, 5mM EDTA, 1.0% SDS, and proteinase inhibitor cocktail from Roche) and sonicated. Protein concentration was determined by using the Bicinchoninic acid assay (BCA, Sigma BCA1-1KT). 30 µg of protein was separated on 10% SDS-PAGE and transferred to a PVDF membrane (Immobilon-P, Millipore). The membrane was blocked with 5% dry fat-free milk solution in PBS containing 0.1% Tween (PBST) at RT for 1 hour and then incubated with indicated primary antibodies overnight at 4°C. The membrane was washed with PBST for three times and incubated with HRP-conjugated secondary antibody for 1 hour at RT followed by three washes using PBST. Immobilon Western chemiluminescent (Millipore WBKLS0500) kit was used to detect HRP activity of the secondary antibody. The membrane was then imaged using ChemiDoc Touch imaging system (Bio-Rad). Image J software was used for band intensity quantification. Primary antibodies used for this study are as follows: RBFOX2 (1:1000, Abcam, ab57154), TPM1 (1:1000, Cell Signaling, D12H4), α-tubulin (1:20000, Sigma-Aldrich, T6074) and ANT1:Slc25a4 (1/750, Abcam, ab102032).

### Immunofluorescence of mitochondrial proteins

H9C2 cells were seeded on to the Nunc lab-tek chamber slides for overnight and then fixed with 4% paraformaldehyde in PBS for 10 minutes followed by permeabilization and blocking in 0.1% triton X100 in 10% goat serum for 2 hours. Cells were then incubated overnight with rabbit anti-OPA1 (Abcam ab157457) and mouse anti-MFN1 (Abcam ab57602) monoclonal antibodies, diluted in 10% goat serum. Then, cells were washed with PBS, incubated with Alexa Flour 594-conjugated goat anti-mouse or Alexa Flour 488-conjugated goat anti-rabbit secondary antibody, and slides were mounted using VECTASHIELD Antifade Mounting Medium with DAPI (Vector Laboratories, Burlingame, CA). Slides were imaged using Zeiss LSM 880 with Airyscan confocal microscope with 63X 1.4 oil immersion objective and images were processed using Image J software.

### Transmission Electron Microscopy

To obtain ultrastructural analysis of cells in ultrathin sections, cells were fixed for at least 1 hour in a mixture of 2.5% formaldehyde prepared from paraformaldehyde powder, and 0.1% glutaraldehyde in 0.05M cacodylate buffer pH 7.3 to which 0.01% picric acid and 0.03% CaCl_2_ were added. Cell monolayers were washed in 0.1 M cacodylate buffer, were scraped off and processed further as a pellet. The pellets were post-fixed in 1% OsO_4_ in 0.1M cacodylate buffer pH 7.3 for 1 hour, washed with distilled water and *en bloc* stained with 2% aqueous uranyl acetate for 20 min at 60°C. The cell pellets were dehydrated in ethanol, processed through propylene oxide and embedded in Poly/Bed 812 (Polysciences, Warrington, PA). Ultrathin sections were cut on Leica EM UC7 ultramicrotome (Leica Microsystems, Buffalo Grove, IL), stained with lead citrate and examined in a JEM-1400 (JEOL USA, Peabody, MA) transmission electron microscope at 80 kV. Digital images were acquired with a bottom-mounted CCD camera Orius SC200 1 (Gatan, Pleasanton, CA).

### Luciferase activity of *Slc25a4* 3**’**UTR constructs

The 3’end of rat *Slc25a4* 3’UTR sequence, which contains multiple poly(A) sites (pPAS and pPAS) and RBFOX2 binding sites downstream of dPAS were cloned into pmirGLO Dual-luciferase vector (Promega, USA) downstream of firefly luciferase open reading frame (ORF). SV40 polyA site was removed in this plasmid to allow only the usage of polyA sites provided within the *Slc25a4* 3’UTR. Two RBFOX2 binding motifs near the distal poly(A) sites in the 3’UTR of *Slc25a4* gene were abolished by mutating “CA” to “AC” (*Slc25a4* 3’UTR RBFOX2 binding WT or *Slc25a4* 3’UTR RBFOX2 binding MUT). 1X10^6^ HEK293 cells seeded in 6 well plates 18h before transfection, and transfected with 750 ng DNA (Luciferase *Slc25a4* 3’UTR polyA reporter constructs) using X-tremeGene 9 (Sigma Aldrich, USA). Cells were lysed 24 hours after transfection and determined by the Dual luciferase assay kit (Promega, USA) and recorded using BioTek Cytation 5 plate reader. Firefly luciferase was normalized to renilla luciferase activity and compared between WT and mutant luciferase *Slc25a4* 3’UTR polyA reporter constructs.

